# An evolutionarily conserved Lhx2-Ldb1 interaction regulates the acquisition of hippocampal cell fate and regional identity

**DOI:** 10.1101/2020.08.24.265207

**Authors:** Veena Kinare, Archana Iyer, Hari Padmanabhan, Geeta Godbole, Tooba Khan, Zeba Khatri, Upasana Maheshwari, Bhavana Muralidharan, Shubha Tole

## Abstract

Protein cofactor Ldb1 regulates cell fate specification by interacting with LIM-homeodomain (LIM-HD) proteins in a tetrameric complex consisting of an LDB:LDB dimer that bridges two LIM-HD molecules, a mechanism first demonstrated in the *Drosophila* wing disc. Here, we demonstrate conservation of this interaction in the regulation of mammalian hippocampal development, which is profoundly defective upon loss of either *Lhx2* or *Ldb1*. Electroporation of a chimeric construct that encodes the Lhx2-HD and Ldb1-DD (dimerization domain) in a single transcript cell-autonomously rescues a comprehensive range of hippocampal deficits in the mouse *Ldb1* mutant, including the acquisition of field-specific molecular identity and the regulation of the neuron-glia cell fate switch. This demonstrates that the LHX:LDB complex is an evolutionarily conserved molecular regulatory device that controls complex aspects of regional cell identity in the developing brain.

**Summary statement:** Similar to an Apterous-Chip mechanism that patterns the Drosophila wing blade, interaction between mammalian orthologs Lhx2 and Ldb1 regulates multiple aspects of hippocampal development in the mouse.

## Introduction

Transcription factors act in macromolecular complexes comprising multiple interacting factors wherein individual components play unique structural and biochemical roles to locally modify the chromatin landscape. While great progress has been made in identifying individual molecules that participate in a given complex, few complexes have been directly tested for identified molecular interactions and functional consequences. One elegant and well-characterized molecular assembly is the “tetrameric model” for LIM-homeodomain (LIM-HD) transcription factor function, which was first demonstrated for LIM-HD protein Apterous (Ap) in the context of its role as a “dorsal selector” in the *Drosophila* wing blade (van Meyel *et al.*, 1999). This study demonstrated that it is necessary for Ap to bind with cofactor Chip/Ldb1 and for Chip to dimerize, thus bringing together two Ap molecules. This active tetramer could be replaced by a chimeric dimer that contained the HD of Ap and the dimerization domain (DD) of Chip, and this was sufficient to restore the normal patterning of the wing blade (Milan and Cohen, 1999; van Meyel *et al.*, 1999).

The interaction between LDB and LIM-HD proteins seen in *Drosophila* is evolutionarily conserved (Thaler *et al.*, 2002), but has thus far not been examined in the mammalian brain. The vertebrate ortholog of *Ap*, *Lhx2,* is the only LIM-HD gene to be expressed in the proliferating progenitors of the entire cortical neuroepithelium from the earliest stages of corticogenesis (Retaux *et al.*, 1999; Bulchand *et al.*, 2001). Lhx2 functions as a “cortical selector,” such that constitutive loss of Lhx2 causes loss of the entire cortical primordium and expansion of two flanking structures, the hem and the antihem (Bulchand *et al.*, 2001; Mangale *et al.*, 2008). In mosaic experiments, patches of *Lhx2* null cells interspersed in wild-type medial cortical primordium differentiate into hem (Mangale *et al.*, 2008). Each patch of hem functions as a “hippocampal organizer,” and induces hippocampal fate in adjacent Lhx2-expressing tissue (Mangale *et al.*, 2008; Godbole *et al.*, 2018). These studies demonstrated that Lhx2 represses hem fate in the medial telencephalon, and consistent with this, the wild-type embryonic hippocampal primordium expresses a high level of *Lhx2* whereas the adjacent hem shows no detectable expression (Bulchand *et al.*, 2001). This raises an important open question as to whether the requirement for Lhx2 in the hippocampal primordium is limited to preventing this tissue from itself differentiating into hem. It remains untested whether Lhx2 functions within presumptive hippocampal cells to execute the instructive cues and direct hippocampal fate in this tissue. At later embryonic stages, when hippocampal neurogenesis is underway, conditional loss of Lhx2 in hippocampal progenitors induces premature astrogliogenesis (Subramanian *et al.*, 2011). The molecular mechanism of Lhx2 function in this process remains unclear and the tetrameric model has never been examined with respect to these functions.

In vertebrates, *Chip* has two homologs, both of which are expressed in the developing mammalian cerebral cortex. Of these, *Ldb1* is ubiquitously expressed, and *Ldb2* has a limited expression and is absent from the telencephalic ventricular zone where progenitors reside (Bach *et al.*, 1997; Bulchand, Subramanian and Tole, 2003). Cortex-specific loss of *Ldb1* results in an extremely shrunken hippocampus, whereas loss of *Ldb2* has no detectable effect on hippocampal development (Leone *et al.*, 2017). In the current study, we found that loss of *Ldb1* from embryonic day (E) 10.5 results in a profound loss of the hippocampal primordium from the earliest stages of development, and also an upregulation of GFAP, suggesting enhanced gliogenesis. We tested whether an Ldb1:Lhx2 complex is critical for these processes by designing a chimeric molecule that comprises the HD of Lhx2 and the DD of Ldb1, modeling our construct based on the seminal studies performed in the *Drosophila* wing and the vertebrate spinal cord (van Meyel *et al.*, 1999; Thaler *et al.*, 2002). Using *in utero* electroporation, we demonstrate that this chimeric molecule not only rescues the appearance of hippocampal tissue that expresses field-specific markers, but is also sufficient to suppress astrogliogenesis and promote neurogenesis in an *Ldb1* loss of function background. While Ldb1 binding partners are not limited to the LIM-HD family (Ramain *et al.*, 2000; Bronstein *et al.*, 2010), this construct selectively restores Lhx2-Ldb1 function in the *Ldb1* mutant. Our results demonstrate an elegant conservation of the Lhx2:Ldb1 molecular toolkit that is required for fundamental aspects of development in the brain, and provides a function for the intense expression of both genes in the early hippocampal primordium.

## Results and Discussion

*Lhx2* and *Ldb1* are both expressed in the dorsal telencephalon at E10.5 in neuroepithelium that will form the hippocampus and neocortex (Figure 1A, B). *Lhx2* expression excludes the hem, which forms at the telencephalic midline (arrowhead, Figure 1A). This pattern of expression is maintained through subsequent stages, such that at E12.5, the entire dorsal telencephalic neuroepithelium except the hem expresses *Lhx2* (Bulchand, Subramanian and Tole, 2003), and this expression persists upon cortex-specific deletion of *Ldb1* using Emx1Cre (Figure 1B, Supplementary Figure 1). Within this *Lhx2*-expressing territory, the neuroepithelium of the hippocampal primordium expresses *EphB1* in control brains (white asterisk, Figure 1B). This expression is greatly reduced or undetectable when either *Lhx2* or *Ldb1* are deleted using Emx1Cre (arrowheads, Figure 1B). Since the hippocampus does not form in the absence of Wnt signaling from the hem (Lee *et al.*, 2000), we examined the expression of *Wnt3a* and *Wnt2b* in *Emx1Cre;Ldb1*^*lox/lox*^ embryos. The expression of both these Wnt genes was apparently normal in the hem of *Ldb1* mutant brains at E12.5 (Figure 1C). Therefore, the near-complete lack of specification of the hippocampal primordium in the *Ldb1* conditional mutants could not be attributed either to the lack of Wnt gene expression in the hem, or lack of *Lhx2* expression in the hippocampal primordium.

**Figure 1:**
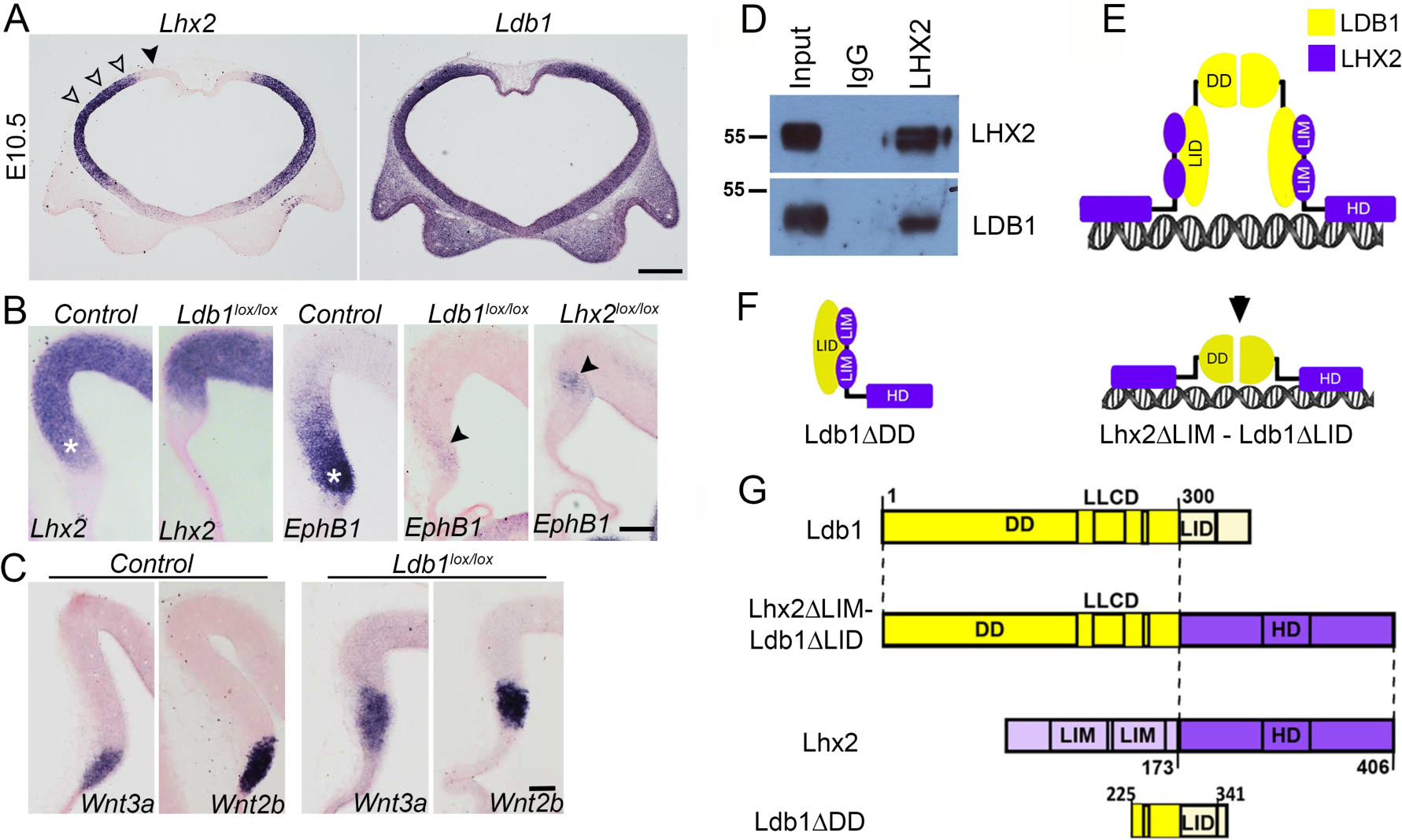
*Ldb1* and *Lhx2* are each required for specification of the hippocampal primordium. (A) Sections from E10.5 wild-type brains display *Lhx2*(A) and *Ldb1*(B) expression in the cortical primordium (open arrowheads). *Lhx2* is undetectable in the hem (arrowhead). (B,C) Sections from E12.5 control and *Emx1Cre; Ldb1*^*lox/lox*^ or *Emx1Cre; Lhx2*^*lox/lox*^ embryos. *Lhx2* expression persists upon loss of Ldb1. *EphB1* expression is severely reduced upon loss of either Ldb1 or Lhx2 (arrowheads). White asterisks mark the hippocampal primordium (B). *Wnt3a* and *Wnt2b* expression appears unaffected in the *Ldb1* mutant hem (C). (D) Immunoprecipitation with anti-Lhx2 antibody pulls down Ldb1 from hippocampal lysate. (E-G) The tetrameric model of LIM-HD (Lhx2/AP) and LDB (Ldb1/Chip) protein interaction and design of the chimeric construct Lhx2ΔLIM-Ldb1ΔLID and Ldb1ΔDD (Milan and Cohen, 1999; van Meyel *et al.*, 1999). Scale bars: 200 μm (A); 100 μm (B,C).

The lack of hippocampal specification in both the *Ldb1* and *Lhx2* mutants, together with the ability of Lhx2 to bind to Ldb1 *in vitro* (Agulnick *et al.*, 1996) and *in vivo* (Gueta *et al.*, 2016; de Melo *et al.*, 2018) is consistent with the well-established tetrameric model of LIM-HD (Lhx2/Ap) and LDB (Ldb1/Chip) protein interaction (Milan and Cohen, 1999; van Meyel *et al.*, 1999) illustrated in Figure 1E. Immunoprecipitation using anti-Lhx2 antibody pulls down Ldb1 and *vice versa*, (Gueta *et al.*, 2016; de Melo *et al.*, 2018). Anti-Lhx2 antibody similarly pulls down Ldb1 from E15 hippocampal tissue (Figure 1D).

Two constructs were used in (Milan and Cohen, 1999; van Meyel *et al.*, 1999), to provide key data for the tetrameric model, and we used the mouse counterparts of these same constructs to test Lhx2:Ldb1 interactions in the hippocampus. The first was a truncated Ldb1 construct lacking the dimerization domain, which is capable of sequestering Lhx2 (Ldb1ΔDD; Figure 1F). For the present study, we designed a second construct similar to the chimeric construct used in (Milan and Cohen, 1999; van Meyel *et al.*, 1999). This Lhx2ΔLIM-Ldb1ΔLID construct encodes a protein in which the LIM domains of Lhx2 are deleted, and the remaining Lhx2-HD containing fragment is fused to a portion of the Ldb1 molecule containing the Ldb1-dimerization domain (Figure 1F,G). We tested whether this chimeric construct can rescue the severely shrunken hippocampus in the cortex-specific *Ldb1* conditional mutant (*Emx1Cre; Ldb1*^*lox/lox*^).

The chimeric construct was electroporated into the medial telencephalon of *Emx1Cre; Ldb1*^*lox/lox*^ embryos at E13-E14, when hippocampal neurogenesis is underway. The embryos and the brains were examined at postnatal day 0 (P0; Figure 2). At this stage, pan-hippocampal marker *Zbtb20*, as well as hippocampal field-specific markers *KA1* (field CA3) and *NeuroD* (CA3 and DG), are robustly expressed in control brains (Figure 2A). In contrast, the non-electroporated hemisphere revealed barely detectable expression and that too in an extremely reduced hippocampus (Figure 2B). The electroporated half of the same sections displayed a somewhat larger hippocampus that expressed each marker in a patchy manner (Figure 2C,D). When compared with the corresponding GFP fluorescence image of the same region, it was apparent that the patches of rescued expression indeed correspond well with regions that had been electroporated with the chimeric construct (Figure 2 D-F).

**Figure 2:**
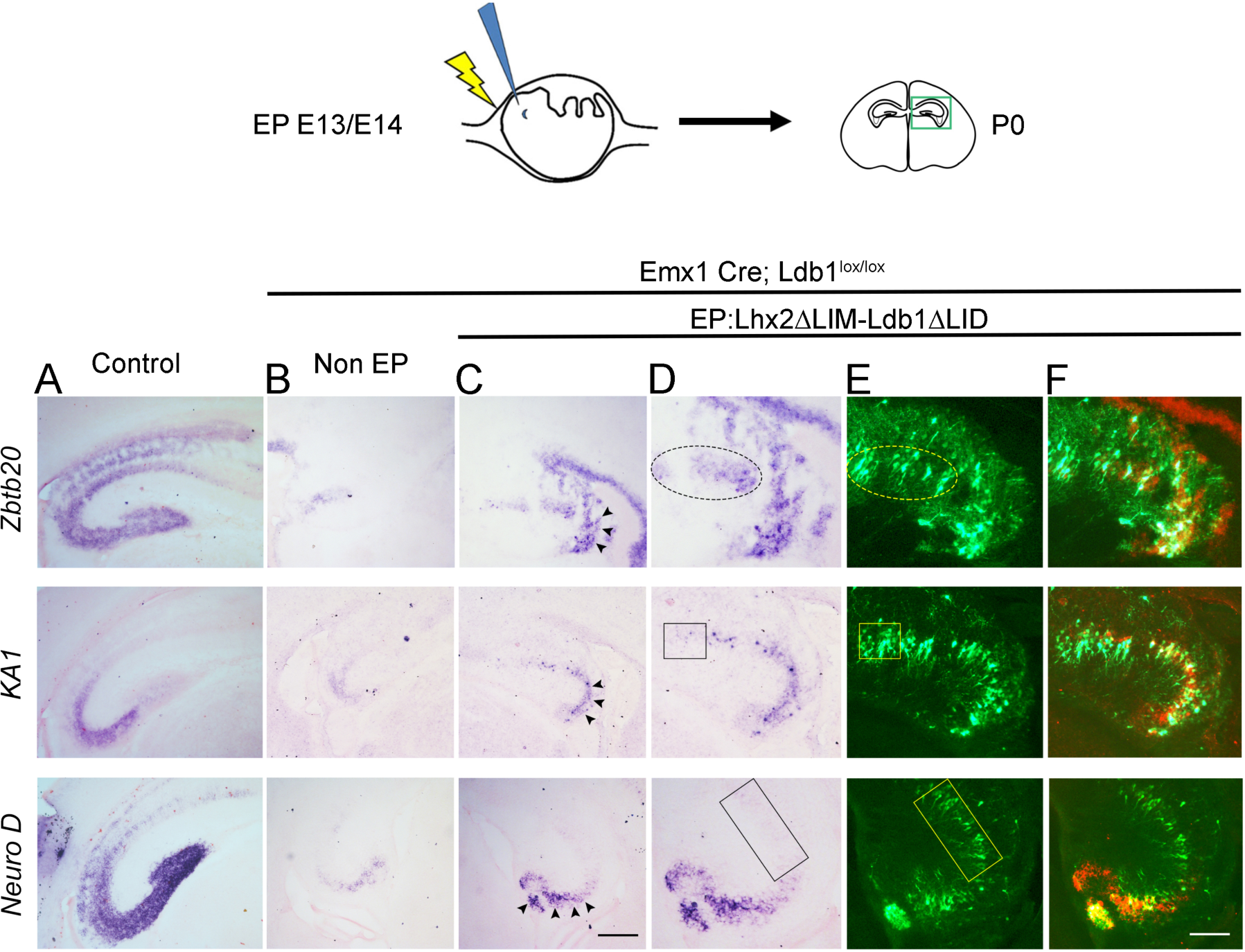
The chimeric construct Lhx2ΔLIM-Ldb1ΔLID rescues hippocampal field-specific marker expression in the absence of *Ldb1*. (A) Sections of control brains at P0 display *Zbtb20* expression in the entire Ammon’s horn and dentate gyrus. *KA1* expression is limited to field CA1, and *NeuroD* expression marks CA3 and the dentate gyrus. (B) Upon loss of *Ldb1*, the expression of each of these markers is greatly reduced. (C) The contralateral hemispheres of the sections in (B), that were electroporated with the chimeric Lhx2ΔLIM-Ldb1ΔLID construct at E13-E14, display a greater intensity and extent of *Zbtb20*, *KA1*, and *NeuroD* expression (arrowheads, C). (D-E) High magnification images of the sections in (C) showing the expression of each marker (D) and the corresponding GFP fluorescence of the same section indicating the extent of electroporation (E). *Zbtb20* expression (dashed oval, D,E), but not *KA1* and *NeuroD* expression (boxes, D, E) is upregulated in electroporated cells at the CA1 end of the Ammon’s horn in *Ldb1* mutant brains. (F) A false-color overlay of the images in (D, E) reveals that the expression of *Zbtb20, KA1*, and *NeuroD* is localized to electroporated regions and does not arise ectopically. Scale bars: 200 μm (A-C); 100 μm (D-F).

It is noteworthy that multiple region- and field-specific features of the hippocampus, lost in the absence of Ldb1, are rescued in a region-appropriate manner by the chimeric construct. Whereas expression of the pan-hippocampal marker *Zbtb20* appears throughout the extent of the electroporated region (ovals in Figure 2 D,E), *NeuroD* and *KA1* expression remains undetectable at the CA1 end even though electroporated cells are present (compare boxes in corresponding images of D, E). Therefore, the introduction of the chimeric construct does not induce widespread upregulation of hippocampal regional markers, but rather, it appears to allow the electroporated cells to read out specification cues in a region-specific manner. This reinforces the interpretation that the chimeric construct substitutes for an endogenous mechanism rendered inoperable by the loss of *Ldb1*.

Previously, we demonstrated that Lhx2 regulates the neuron-glia cell fate switch in hippocampal progenitors. Loss of *Lhx2* during hippocampal neurogenesis causes premature astrogliogenesis, and overexpression of *Lhx2* during the period of astrogliogenesis prolongs neurogenesis (Subramanian *et al.*, 2011). Since the chimeric Lhx2ΔLIM-Ldb1ΔLID construct is capable of restoring the formation of cells with specific hippocampal field identities, we further sought to test if it is also capable of restoring neurogenesis in the absence of either Lhx2 or Ldb1.

We used the dominant-negative construct, Ldb1ΔDD, in which the dimerization domain of *Ldb1* was deleted (Figure 1H), to sequester Lhx2 in E14.5 hippocampal progenitors. Previously, we showed that the protein encoded by this construct does indeed bind the LIM-domains of Lhx2 (Ldb1ΔDD is called ClimΔDD; Supplementary Figure S1(Subramanian *et al.*, 2011). We further demonstrated that electroporation of this construct causes a premature gliogenic effect indistinguishable from that seen upon electroporation of Cre-GFP into *Lhx2*^*lox/lox*^ embryos (Supplementary Figure S2; (Subramanian *et al.*, 2011). Thus, Ldb1ΔDD recapitulates the functions of its *Drosophila* counterpart ChipΔDD, which binds and sequesters Apterous, but cannot form a functionally active tetrameric complex (Bach *et al.*, 1999; van Meyel *et al.*, 1999; Becker *et al.*, 2002). The Drosophila counterpart of the chimeric Lhx2ΔLIM-Ldb1ΔLID construct also effectively substitutes for loss of Ap and rescues *Ap* mutant phenotypes in Drosophila (van Meyel *et al.*, 1999). We therefore used Ldb1ΔDD and the chimeric Lhx2ΔLIM-Ldb1ΔLID construct to further test the tetrameric model in the mouse hippocampus.

In our previous studies, we used GFAP as a marker of astrocytes derived from the hippocampal primordium, and showed that E15 hippocampal progenitors produce either ß-tubulin-expressing neurons or GFAP-expressing astrocytes in the 5–7-day window after electroporation *in vivo* (Subramanian *et al.*, 2011) and *in vitro* (Muralidharan, Keruzore, *et al.*, 2017). We further demonstrated that the GFAP-expressing cells produced by loss of *Lhx2* also express astrocyte marker AldoC, but do not express oligodendrocyte marker Olig2 (Subramanian *et al.*, 2011). Therefore, in the present study, we proceeded with using GFAP as a validated marker for astrocytes.

We performed *in utero* electroporation in wild-type embryos at E14.5, examined the brains 7 days later at P2, and scored for the expression of GFAP in the electroporated cells (Figure 3B). Electroporation of a control GFP construct revealed a baseline gliogenesis of 27%. Electroporation of the chimeric construct alone resulted in 20% gliogenesis, not significantly different from the baseline, which was expected since endogenous *Lhx2* expression is already very high at E14.5 (Bulchand, Subramanian and Tole, 2003; Subramanian *et al.*, 2011) and hence a construct designed to mimic Lhx2 function would not be expected to have a significant additive effect. The dominant-negative Ldb1ΔDD construct caused significantly enhanced gliogenesis (61%), as seen in (Subramanian *et al.*, 2011). This was suppressed by co-electroporation of the chimeric construct to 32%, similar to baseline levels, thus effectively restoring the percentage gliogenesis to control levels (Figure 3C). These results suggest that the chimeric Lhx2ΔLIM-Ldb1ΔLID construct can preserve Lhx2-like functionality even when Lhx2 is sequestered.

**Figure 3:**
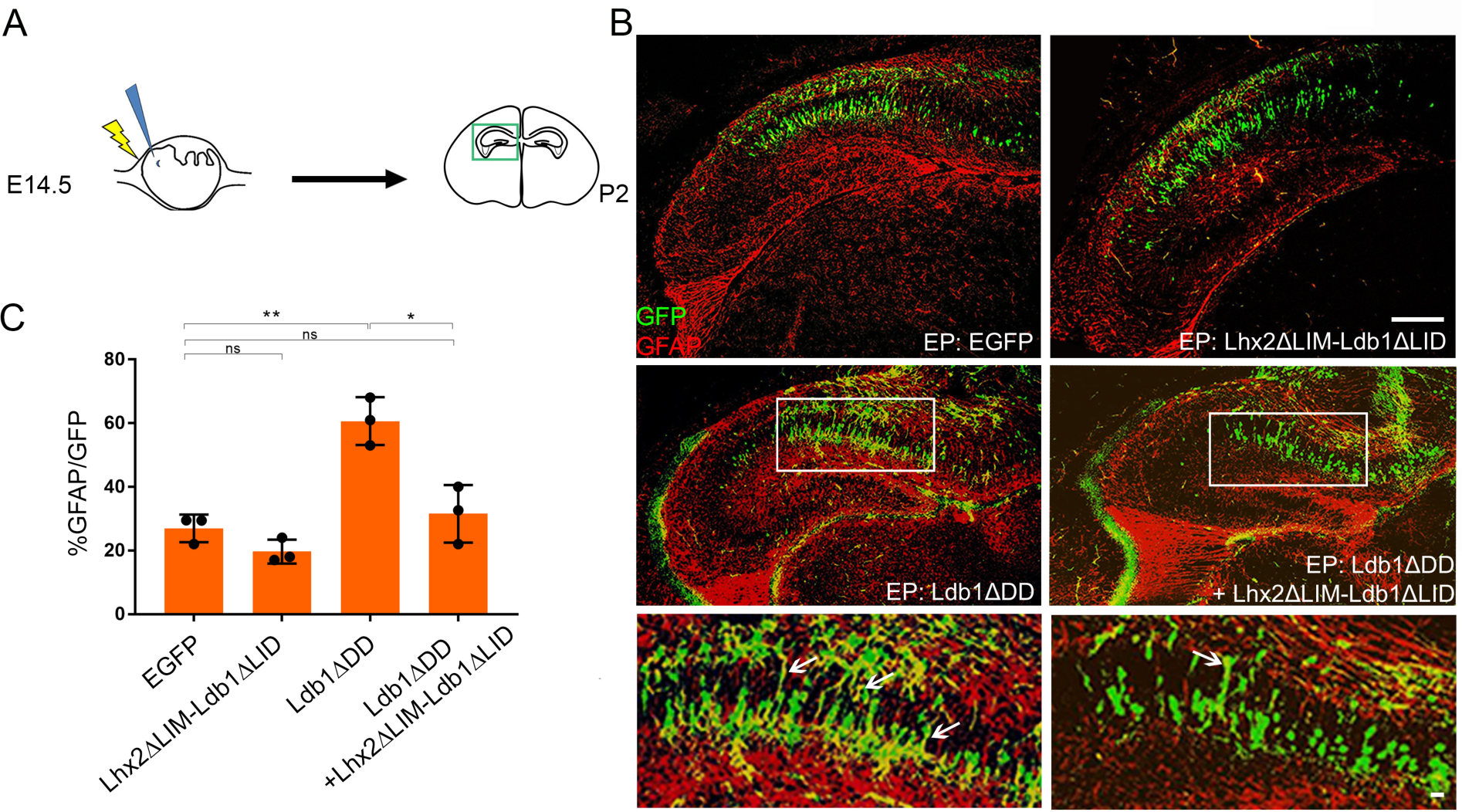
The chimeric construct Lhx2ΔLIM-Ldb1ΔLID suppresses the enhanced gliogenic effect of the dominant-negative Ldb1ΔDD. (A) Cartoon illustrating *in utero* electroporation at E14.5 in wild-type embryos and examination of the hippocampus at P2. (B, C) Electroporation of control GFP or the chimeric construct Lhx2ΔLIM-Ldb1ΔLID produces 27% or 20% GFP(green)-GFAP(red) co-expressing cells, respectively. Electroporation of the dominant-negative Ldb1ΔDD increases this fraction to 61%, but co-electroporation of the chimeric construct restores it to 32%, similar to control levels (arrows indicate GFP(green)-GFAP(red) co-expressing cells). n=3 per condition. Error bars indicate SD. *p<0.05, **p<0.005. Scale bars: 200 μm (low magnification); 10 μm (high magnification) images in B.

Loss of Lhx2 causes premature astrogliogenesis and hence upregulation of GFAP expression in the hippocampus (Subramanian *et al.*, 2011). We examined GFAP expression in *Emx1Cre; Ldb1*^*lox/lox*^ brains at P0, and found apparently enhanced GFAP expression in the rudimentary hippocampus (Figure 4A,B). We established a baseline for the percentage astrogliogenesis at E14.5 in these mutant brains by electroporating GFP at this age and scoring the fraction of GFP+GFAP co-expressing cells at P0. Upon electroporation of a control GFP construct, 43% of the GFP-positive cells in the shrunken *Ldb1* mutant hippocampus were seen to express GFAP (Figure 4C, D). Electroporation of the chimeric construct suppressed astrogliogenesis to 6% (Figure 4D-F). Furthermore, electroporated cells were seen to produce axons that reach the rostral level of the hippocampal commissure, confirming that these cells are indeed neurons (arrowhead, Figure 4F). Together, our results confirm a rescue of hippocampal neurogenesis and a suppression of astrogliogenesis in the cortex-specific *Ldb1* mutant. In summary, these results demonstrate that the chimeric Lhx2ΔLIM-Ldb1ΔLID construct restores hippocampal neurogenesis in the absence of Ldb1.

**Figure 4:**
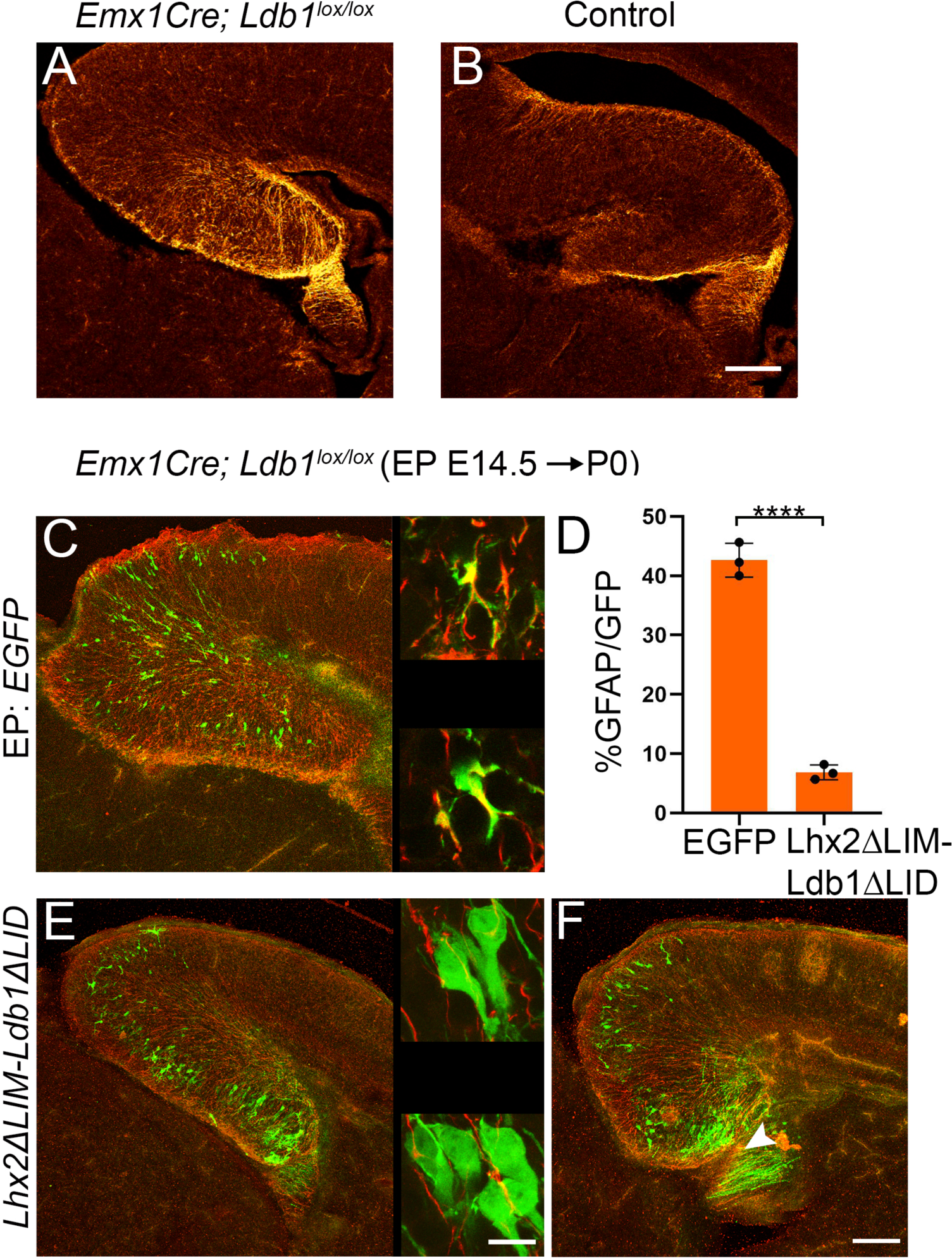
The chimeric Lhx2ΔLIM-Ldb1ΔLID construct suppresses gliogenesis and rescues neurogenesis in the cortex-specific *Ldb1* mutant. (A, B) GFAP immunohistochemistry at P0 in the cortex-specific *Ldb1* mutant hippocampus (A) appears enhanced compared with that seen in wild-type control brains (B). (C-F) Control GFP (C) or the chimeric construct (E,F) was electroporated into cortex-specific *Ldb1* mutants at E14.5 and the hippocampus was examined at P0. High magnification images of cells from C and E are in the corresponding insets. (D) The percentage of GFP-expressing cells(green) that are also GFAP-expressing(red) was scored. The percentage of astrocytes generated from electroporated cells in the cortex-specific *Ldb1* mutant decreases from 43% (control GFP) to 6% (chimeric construct). n=3; Error bars indicate SD. ****p<0.0005. Arrowhead (F) indicates hippocampal axons arising from cells electroporated with the chimeric construct. Scale bar: 200 μm (all low magnification images); 10 μm (high magnification insets in C and E).

## Conclusion

The present study was prompted by the striking similarity in the hippocampal phenotype upon cortex-specific deletion of *Ldb1* and *Lhx2*. In both mutants, the medial telencephalic neuroepithelium corresponding to the hippocampal primordium is shrunken, and displays limited expression of hippocampal markers, indicating a loss or lack of specification of this structure. The central finding of our study is that the introduction of a chimeric construct containing the Lhx2-HD and the Ldb1-DD is sufficient to restore field specification as well as neurogenesis in the developing hippocampus. These findings take on a particular significance in light of the fact that *Lhx2* expression is not lacking in the *Ldb1* mutants, supporting the interpretation that the transcription factor is unable to function in the absence of its cofactor Ldb1. Further studies will seek to identify other members of the multi-protein complex that are likely critical for Lhx2-Ldb1 function (Muralidharan, Khatri, *et al.*, 2017), taking advantage of the fact that the chimeric construct, in effectively substituting for both Lhx2 and Ldb1, may help narrow down a limited set of key binding partners.

*Lhx2* and *Ldb1* are normally expressed in the entire hippocampal progenitor zone, but field-specific features arise in distinct portions of the hippocampal primordium, consistent with the interpretation that the electroporated progenitors may be responding to a morphogen gradient from the hippocampal organizer, the hem, which we found to express apparently normal patterns of *Wnt2b and 3a* in the cortex-specific *Ldb1* mutant. Together, the data support the interpretation that the Lhx2:Ldb1 complex does not alone impart field identity in the hippocampus, but functions to execute the inductive influence of the hippocampal organizer. This extends the established roles of Lhx2 and Ldb1 function.

The fact that Lhx2, a mammalian ortholog of Ap, functions in a molecular complex with Ldb1 in the brain in a manner that is indistinguishable from what was established for Ap function in the *Drosophila* wing (Milan and Cohen, 1999; van Meyel *et al.*, 1999), indicates an exquisite conservation of this evolutionarily ancient mechanism in the developing hippocampus. These results underscore the importance of this particular protein complex as a fundamental toolkit that has been reutilized in diverse organisms and systems ranging from invertebrates to mammals.

## Material and Methods

### Mice

All animal protocols were approved by the Institutional Animal Ethics Committee (Tata Institute of Fundamental Research, Mumbai, India) according to regulations devised by the Committee for the Purpose of Control and Supervision of Experiments on Animals (CPCSEA), India. The mouse lines used in this study were kind gifts from Yangu Zhao, NIH (*Ldb1*^*lox/lox*^; (Zhao *et al.*, 2007), Edwin Monuki, University of California, Irvine, USA (*Lhx2*^*lox/lox*^; (Mangale *et al.*, 2008). The Emx1Cre line was from Jackson Laboratories (strain name: B6.Cg-Emx1tm1(cre)Krj/J; stock number: 005628). Noon of the day of the vaginal plug was designated as embryonic day 0.5 (E0.5). Control embryos used were littermates that did not have the Cre allele.

### *In utero* electroporation, *in situ* hybridization, and immunohistochemistry

These procedures were performed as described in (Subramanian *et al.*, 2011). For the experiments in Figure 3 and 4, sections were immunostained using anti-GFP antibody (Abcam, catalog no. ab6658) and anti-GFAP antibody (Sigma, catalog no. G9269). Probes used were kind gifts from Forbes D. Porter (Lhx2), Elizabeth Grove (EphB1, NeuroD), Jim Boulter (KA1), Clifton Ragsdale (Wnt3a, Wnt2b), Jakob Nielsen (Zbtb20) and Yangu Zhao (Ldb1). All *in situ* hybridization experiments shown in Figure 1 were performed in three embryos of each genotype. For the data in Figures 2–4, three embryos were examined for each condition.

### Immunoprecipitation

This was performed using E15.5 hippocampal tissue lysate, anti-Lhx2 antibody (Santa Cruz, SC 19344) and biotinylated anti-Ldb1 antibody (Santa Cruz, SC 11198).

### Real time PCR

Medial tissue from control and Emx1 Cre; Ldb1^lox/lox^ was harvested at E12.5 and RNA was extracted using RNeasy Micro Kit (Qiagen, Catalog no.74004). cDNA was prepared using SuperScript IV First-strand Synthesis System (Invitrogen, Catalog no. 18091050) followed by real time quantitative PCR using Kapa SYBR Fast qPCR kit (Kapa Biosystems, Catalog No. KR0389_S) and Light Cycler 96 (Roche). The following primers were used to assess Lhx2 expression:

Primer Set1: Forward Primer: 5’ TCTGACCGCTACTACCTGCT-3’, Reverse Primer – 5’ GGGAGGGGCTGTAGTAGTCT-3’

Primer Set 2: Forward Primer: 5’ GCTGAACACCTGGATCGTGA-3’, Reverse primer – 5’ACCAGACCTGGAGGACTCTC-3’

### DNA constructs

The chimeric construct Lhx2ΔLIM-Ldb1ΔLID was cloned into the pEF1-Myc-His A vector. This chimeric construct was designed based on (van Meyel *et al.*, 1999). DNA-encoding amino acids 1-300 of Ldb1 (Ldb1/Clim2 variant3 {NM_010697}) and amino acids 173-406 of Lhx2 were used for the chimeric fusion protein. For electroporation, this pEF1-Lhx2ΔLIM-Ldb1ΔLID-Myc-His chimeric construct was mixed with one encoding pCAG-IRES2-EGFP in a 1:1 ratio for detection of the electroporated cells. The pCAG-Ldb1ΔDD-IRES2-EGFP construct (Ldb1ΔDD/ClimΔDD) was cloned as described in (Subramanian *et al.*, 2011).

### Image Acquisition and Analysis

Confocal images were acquired using the Olympus FV1200, Zeiss 510 and Zeiss LSM 5 Exciter - AxioImager M1 imaging systems. For calculating the percentage gliogenesis, 3 different electroporated brains were scored for each condition. The-numbers of GFP+GFAP double positive cells were expressed as a percentage of the total GFP cells scored in each brain (Subramanian *et al.*, 2011). Statistical analysis was performed using one-way ANOVA (Tukey’s multiple comparison) for Fig 3C. For Fig 4D, unpaired two-tailed Student t-test was used. In Figure 3, the total cells scored per condition were: 290 (GFP), 298 (chimeric construct + GFP), 285 (Ldb1ΔDD) and 305 (chimeric construct + Ldb1ΔDD), and in Figure 4, the total cells scored per condition were: 284 (GFP) and 412 (chimeric construct+GFP).

## Acknowledgements

We thank Edwin S. Monuki for the kind gift of the *Lhx2*^*lox/lox*^ line, and Yangu Zhao, Paul Love, and Sue McConnell for the *Ldb1*^*lox/lox*^ line. We also thank Elizabeth Grove (EphB1, Lhx9, NeuroD), Yangu Zhao (Ldb1), Jim Boulter (KA1), Clifton Ragsdale (Wnt3a, Wnt2b), Jakob Nielsen (Zbtb20) and Forbes D. Porter (Lhx2) for gifts of plasmid DNA; Shital Suryavanshi and the animal house staff of the Tata Institute for Fundamental Research (TIFR) for excellent support; Anindita Sarkar and Lakshmi Subramanian for critical input on the manuscript. We gratefully acknowledge the support and mentorship of Medha Rajadhyaksha (VK). This work was supported by a University Grants Commission fellowship (VK); a DST-INSPIRE faculty fellowship DST/INSPIRE/04/2018/001140 (AI); Wellcome Trust-Department of Biotechnology India Alliance Early Career Fellowships IA/E/11/1/500402 (GG) and 500197/Z/11/Z (BM); grants from the Department of Biotechnology (PR8681) and the Department of Science and Technology (DST/CSRI/2017/202-G), Ministry of Science and Technology, Govt. of India (ST). We acknowledge the support of the Department of Atomic Energy, Government of India, under project no. RTI4003.

