## Supplementary Figure 1 for "An evolutionarily conserved Lhx2-Ldb1 interaction regulates the acquisition of hippocampal cell fate and regional identity"

Supplementary Fig S1:

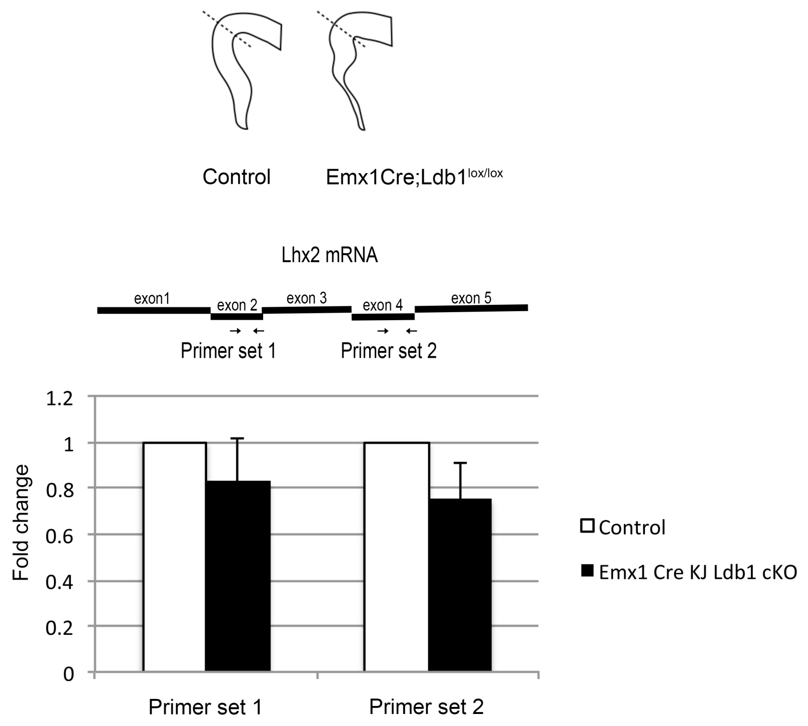


**Supplementary Figure 1: Lhx2 expression levels remain unchanged between control and cortex specific Ldb1 mutant**

Schematic of the tissue dissected for qPCR. Primers used to detect Lhx2 expression. Fold change using qPCR between control and Ldb1 mutant tissue showing no change in Lhx2 expression.
